# Where are they now? Tracking the Mediterranean Lionfish Invasion via Local Dive Centers

**DOI:** 10.1101/2020.12.18.423475

**Authors:** Elizabeth W Phillips, Alexander Kotrschal

## Abstract

Invasive species are globally on the rise due to human-induced environmental change and are often a source of harm to their new ecosystems. Tracking the spread of invaders is crucial to better management of invasive species, and citizen science is often used to collect sighting data. However, this can be unreliable due to the general public’s limited expertise for accurate identification and a lack of clear absence data. Here, we introduce a refined method of citizen science by tracking the spread of the invasive lionfish (*Pterois miles*) in the Mediterranean Sea using dive centers’ expertise on local marine wildlife. We contacted 1131 dive centers on the Mediterranean coast via email and received 216 responses reporting whether or not lionfish were present in their area and, if present, the year they were first sighted. Currently, lionfish sightings are observed in the eastern half of the Mediterranean, though the front is continuing to move west with the furthest sighting as far as Corfu, Greece (19.939423°E, 39.428017°N). In 2020, lionfish also expanded their invasive range north on the Turkish Aegean coast to Karaburun (26.520657°E, 38.637033°N), showing that the invasion is ongoing. We found that the invasive range is now exceeding previous invasion models, highlighting the need for additional research on lionfish biology to inform management efforts. Continuous monitoring of invasive fronts based on dive center reports and a better understanding of what makes lionfish so invasive is crucial to mitigating their negative impact on native ecosystems.

## 1. Introduction

As humans continue to shape and modify their environment, animals often find themselves in novel situations to which they must react and adapt. One of these situations is when direct human transport or man-made constructions allow animals to move from one area to the other and become invasive. Globally, invasive species have been on the rise due to human-induced environmental change, and often cause ecological damage to the areas they invade (Cassey, Blackburn, Duncan, & Chown, 2005; Simberloff et al., 2013). A crucial step to creating effective management strategies for invasive species is tracking their introduction and expansion into new regions. Often citizen science databases, like “Is it Alien to you? Share it!!!”or Seawatchers, are used by researchers to gain insight into new invasive species or expansion on the front of existing invasive species (Dimitriadis et al., 2020; Katsanevakis et al., 2020). These databases offer a platform for the general public to report and collect data for professional scientists to use in a collaborative effort, often tracking species sightings across larger areas (Larson et al., 2020). Although these types of reports often allow researchers to gain larger quantities of data than what could be collected alone, citizen scientists may not have the expertise to reliably identify unfamiliar species and usually these reports only list confirmed sightings, so clear absence data is not obtainable (Crall et al., 2010; Larson et al., 2020; Sandahl & Tøttrup, 2020).

Sighting records of invasive lionfish (*Pterois miles*) in the Mediterranean is one example of species tracking using citizen science. In 1869 the Suez Canal opened up a new passageway for marine organisms to move between the Red Sea and the Mediterranean either by currents carrying pelagic larvae or by adult movement through the channel (B. S. Galil et al., 2014). Though the canal is open to movement from either direction, species mainly migrate from the Red Sea to the Mediterranean because of current direction and increased habitat flexibility and biodiversity in Red Sea-native organisms (reviewed in Mavruk & Avsar, 2008). These “Lessepsian species” totaled 63 by 2008 and have been on the rise due to the recent expansion of the Suez Canal (Bella S Galil et al., 2015; Mavruk & Avsar, 2008). One of these species, the lionfish (*Pterois miles*) was rarely sighted in the Mediterranean until 2012 when sightings began to rise and their invasion commenced (Bariche, Torres, & Azzurro, 2013; Golani & Sonin, 1992; Jimenez et al., 2016). With their recent expansion into the Mediterranean, concern has risen about its ecological impacts based on the well-studied lionfish invasion in the Atlantic. Release of lionfish from the aquarium trade in Florida, USA has allowed them to spread throughout the Western Atlantic Ocean and Caribbean Sea with numerous detrimental effects on native communities, including decreases in biodiversity, reef deterioration, and economic loss in the fishing industry (Côté, Green, & Hixon, 2013; Henderson, 2012; Lesser & Slattery, 2011). Their rapid expansion has been attributed to characteristics like their high fecundity, venomous spines, generalist diet, and habitat flexibility (Côté et al., 2013; Galloway & Porter, 2019; Peake et al., 2018; Rojas-Vélez, Tavera, & Acero, 2019). These characteristics are likely to facilitate the lionfish’s invasion in the Mediterranean as well, with similar negative impacts on the Mediterranean ecosystem if allowed to spread (Galanidi, Zenetos, & Bacher, 2018; Kletou, Hall-Spencer, & Kleitou, 2016; Ulman et al., 2020).

Initial reports of the invasion showed lionfish spread and establishment around Cyprus and on the Levantine Coast located at the eastern border of the Mediterranean, but the most recent report documented sightings as far west as the Greek Ionian coast, as well as single sightings of lionfish in Italy and Tunisia (Ernesto Azzurro, Stancanelli, Di Martino, & Bariche, 2017; Dimitriadis et al., 2020). These reports come from small studies of individual countries, or use citizen science databases to track the spread throughout the entire Mediterranean Sea (E. Azzurro & Bariche, 2017; Dimitriadis et al., 2020; Katsanevakis et al., 2020; Kletou et al., 2016; Özbek, Mavruk, Saygu, & Öztürk, 2017). To create a more complete and reliable overview of the lionfish invasion in the Mediterranean, we need to utilize people that have enough local expertise to be able to reliably identify the invasive fish and can say with some confidence that there are no lionfish present if they do not see them. This type of local underwater expertise can be found at dive centers, and centers can be found all over the Mediterranean (Isabel, Martin, Gelcich, Stotz, & Thiel, 2021). We conducted a survey of dive centers around the Mediterranean and asked them to report if they have seen lionfish in their area, and if so, when they first appeared in order to create the most up-to-date report of the lionfish invasion in this area.

## 2. Methods

### 2.1 Dive Center Search

Google Maps was used to locate dive centers situated along the Mediterranean Sea using the search term “dive center”. The search was started at the mouth of the Suez Canal and continued westward along the coast, circling back to the start point, and included all islands located in the sea. Care was taken to ensure all costal area was searched and that all relevant dive center results would be revealed in an area. Inclusion criteria for dive centers contacted for the survey were:

- Appeared in the first 20 results listed for a given area
- Offered tours or courses in diving (ensure active divers on staff)
- Had their own website with contact information listing at least one email (no website contact forms were used)
- Named on Google Maps written using mostly the Latin alphabet

When a dive center was found that met our criteria, their website was searched for all available emails related to diving and recorded, along with the geographic coordinates of the dive center. This yielded 1238 emails from 1131 dive centers. The distribution of the dive centers contacted is shown in Figure 1.

**Figure 1.**
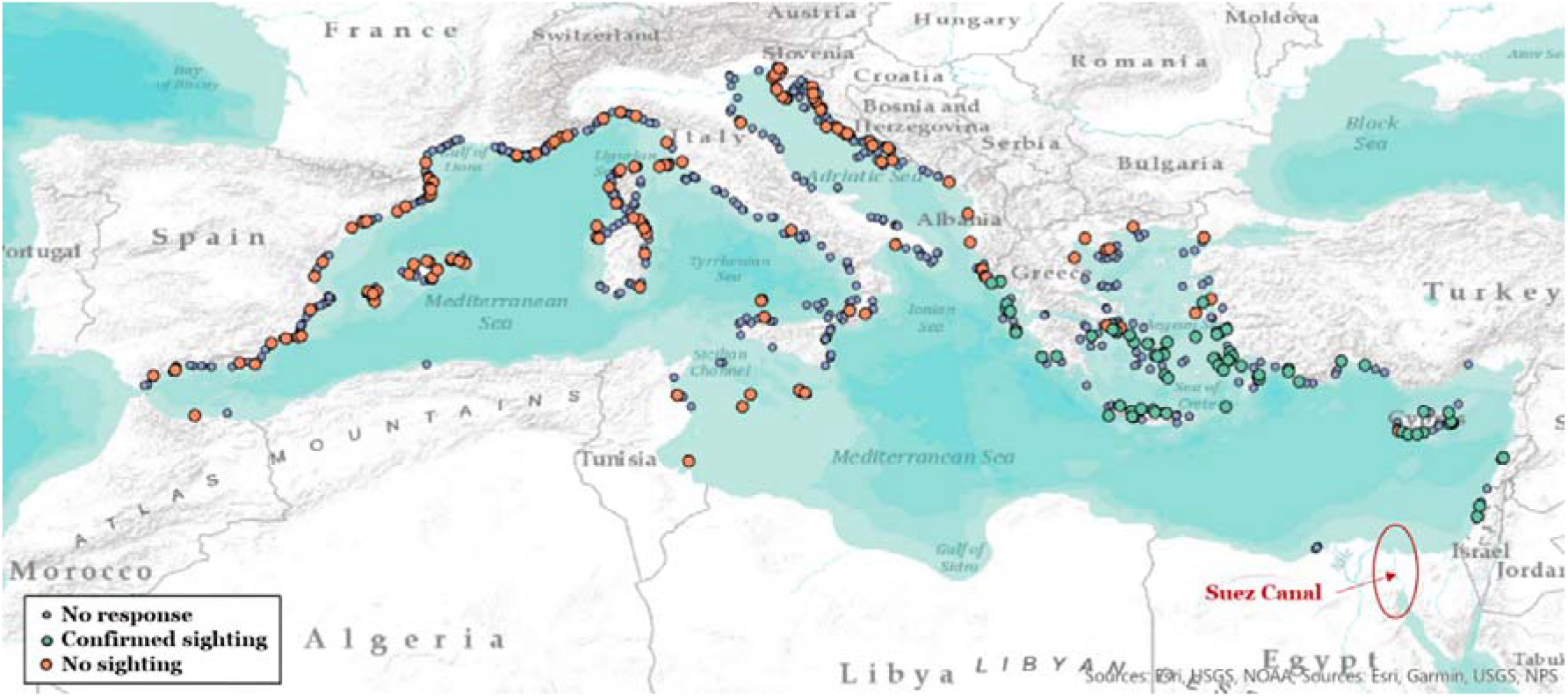
Contacted dive centers and survey respondents. Each dot represents a dive center that was contacted during our survey. Green dots represent survey respondents that reported confirmed sightings of lionfish and red dots are confirmed no-sightings. Gray dots are dive surveys that were contacted, but gave no response. Please note that the one response from Tossa de mar, Spain (2.9030443°E, 41.7109527°N) discussed in the main text is labelled as gray as this sighting may not be robust.

### 2.2 Survey Distribution

We contacted dive centers found through Google Maps via email and asked two questions: 1) Have you seen lionfish in your area? and 2) If so, when did you first see them?. Email text was written in English and translated into 7 other languages (Arabic, French, Spanish, Italian, Turkish, Greek, and German) by native speakers when possible in order to make the survey more accessible to those who do not speak English fluently. If after 1 week no response was received from a dive center, a reminder email was sent in order to maximize the number of responses received.

### 2.3 Data Analysis

Responses from individual dive centers and coordinates previously recorded during the search process, were imported into ArcGIS Pro (version 2.5.1) to create maps illustrating lionfish sightings based on dive center coordinates (Ersi, 2020). We choose to use dive center coordinates as locations of lionfish sightings since coastal dive centers typically use dive sites close to their dive center.

## 3. Results

After contacting 1131 dive centers, we received 216 responses. Of these responses, 75 reported sightings of lionfish in their local area and 141 reported that no lionfish were present (Figure 1). We found that lionfish had been sighted at least once in Israel, Lebanon, Cyprus, Turkey, Greece, and Spain, while dive centers in Tunisia, Morocco, France, Italy, Malta, Slovenia, Croatia, Montenegro, and Albania only reported no sightings. With the exception of one dive center on the south-west coast of Cyprus, dive centers from the east side of the Mediterranean up until the Turkish Aegean coast and mainland Greece unanimously reported confirmed lionfish sightings. In the Aegean Sea, lionfish sightings were reported as far north as Karaburun, Turkey (26.5206573°E 38.6370333°N), and in the Ionian Sea the northern-most sighting, and the western-most sighting overall, was at Corfu Island, Greece (19.9394234°E 39.4280170°N).

For confirmed sighting reports, we also recorded the year that lionfish were first spotted in the area as reported by dive centers. We found that single lionfish sightings were observed in 1997 on a Greek island off the coast of Turkey (27.3126756°E 36.8851393°N), near Tossa de mar, Spain in 2002 (2.9030443°E, 41.7109527°N), Antalya, Turkey in 2005 (30.7054499°E 36.8767709°N), Ölüdeniz, Turkey in 2013 (29.1116429°E, 36.5499014°N) and Yeni Iskele, Cyprus in 2014 (33.949655°E, 35.312925°N). Establishment of lionfish occurred in Cyprus and Turkey by 2015, with the invasion front continuing to move west and north each year (Figure 2). In 2016, lionfish were first spotted near the island of Crete (24.3911364°E 35.1907813°N and 26.6803124°E 35.2259198°N), and in 2018 they were seen off of mainland Greece in the Ionian Sea (21.6503151°E 36.9944446°N). In the Aegean Sea and Ionian Sea, the furthest northern sightings as documented above were in the years 2020 and 2019, respectively.

**Figure 2.**
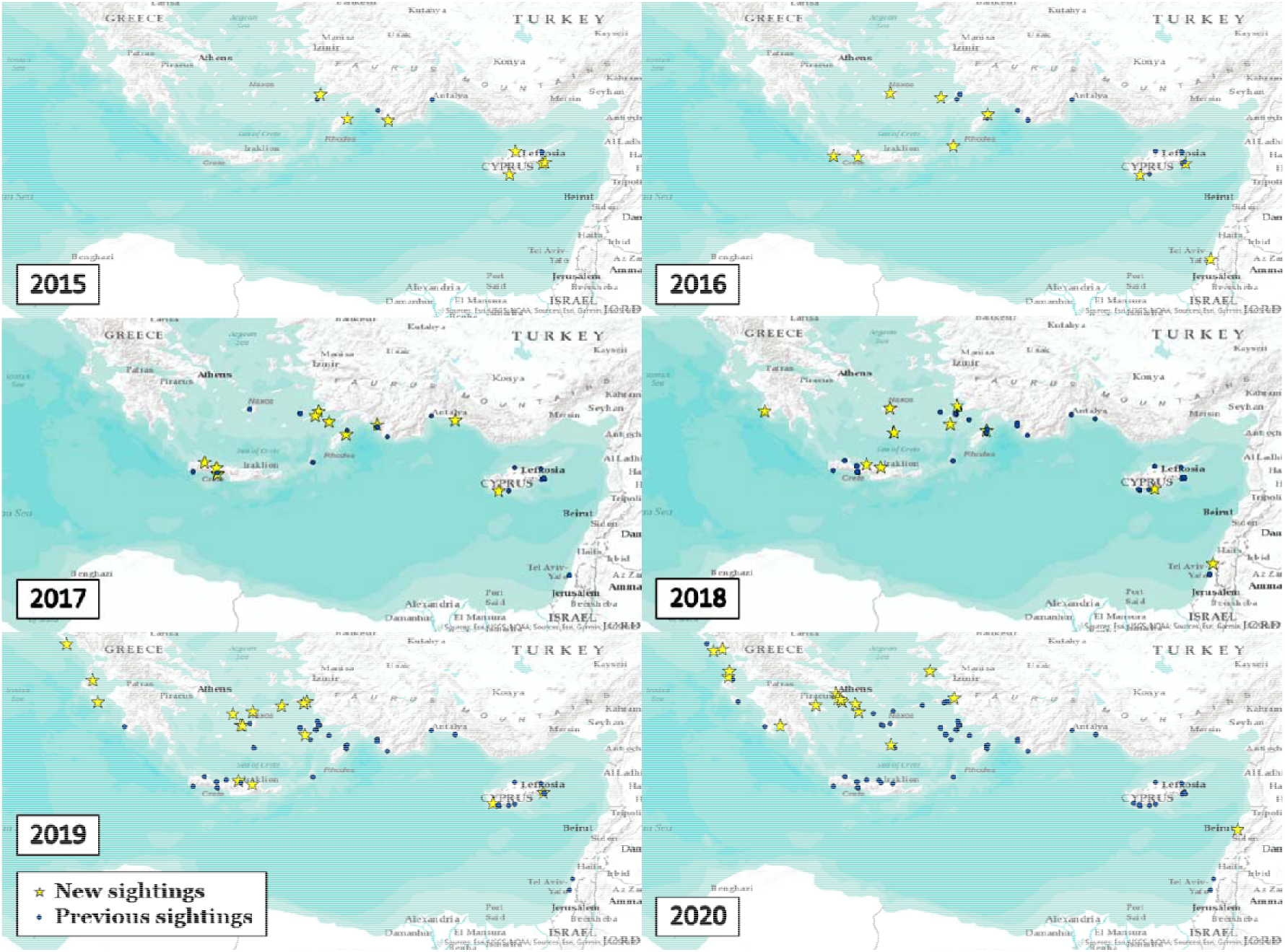
Lionfish sightings by year. Yellow stars represent new sightings of lionfish fish in the given year and blue dots represent older sightings from previous years.

## 4. Discussion

Based on the responses of 216 Mediterranean dive centers, lionfish were observed irregularly around Turkey from 1997-2014, but establishment was first seen in Cyprus and Turkey in 2015 with multiple sightings in these areas. We found a boom of sightings during the following years with a westward movement of the front along Turkey, Crete, and mainland Greece. This is consistent with the invasion spread reported using citizen science and local expertise at a smaller, national scale (Ernesto Azzurro et al., 2017; Dimitriadis et al., 2020). Apart from our novel methodology of tapping into the local expertise of hundreds of professional divers, our study is the first to investigate the most recent spread, including data from 2020, and to document the sighting of lionfish in Spain. Due to the distance between the majority of sightings and that in Spain, we speculate that this animal was either an aquarium release without establishment or a misidentification of another species. Sightings like this are important to note though, as releases from the aquarium trade in places like the Bay of Marseilles, France are predicted to result in high invasion pressure from lionfish (Johnston & Purkis, 2014).

Although we found no further establishment west of Corfu, Greece (reported in 2019), we found an ongoing expansion north in 2020 to Karaburun, Turkey. This movement north in recent years is noteworthy, as a thermal barrier has long been considered a prominent hindrance to lionfish invasion across the entire Mediterranean Sea. In the Atlantic Ocean, lionfish only establish in locations with mean winter temperatures above 15.3°C, though lionfish have experimentally been shown to withstand temperatures as low as 10°C (Kimball, Miller, Whitfield, & Hare, 2004; Whitfield et al., 2014). These isotherm boundaries are currently only present north of the lionfish sightings, though just barely, so it remains to be seen in the coming years if lionfish in the Mediterranean will be able to cross the thermal barrier and establish in more northern regions, or if they will be limited in their spread (Figure 3; Dimitriadis et al., 2020; Johnston & Purkis, 2014). In the Atlantic, lionfish are only seen north of this barrier during summer’s high temperatures, but none have been able to establish year-round. In the next decades, climate change is also predicted to push the isotherms further north, likely allowing lionfish to invade the entire Mediterranean by 2100 (Dimitriadis et al., 2020; Turan, 2020). Importantly, several models based on other biogeographic barriers such as currents predict limited spread of lionfish in the Mediterranean past their present established territories (Johnston & Purkis, 2014; D Poursanidis, 2015; Dimitris Poursanidis, Kalogirou, Azzurro, Parravicini, & Bariche, 2020). Current patterns in the Mediterranean Sea are thought to cause low connectivity between suitable habitats for lionfish, limiting potential larvae spread to the west (Johnston & Purkis, 2014). However, our data shows that lionfish have spread past these assumed invasion boundaries. What other factors facilitate this spread beyond previously suggested boundaries remains unknown Here, we offer several non-mutually exclusive explanations.

**Figure 3.**
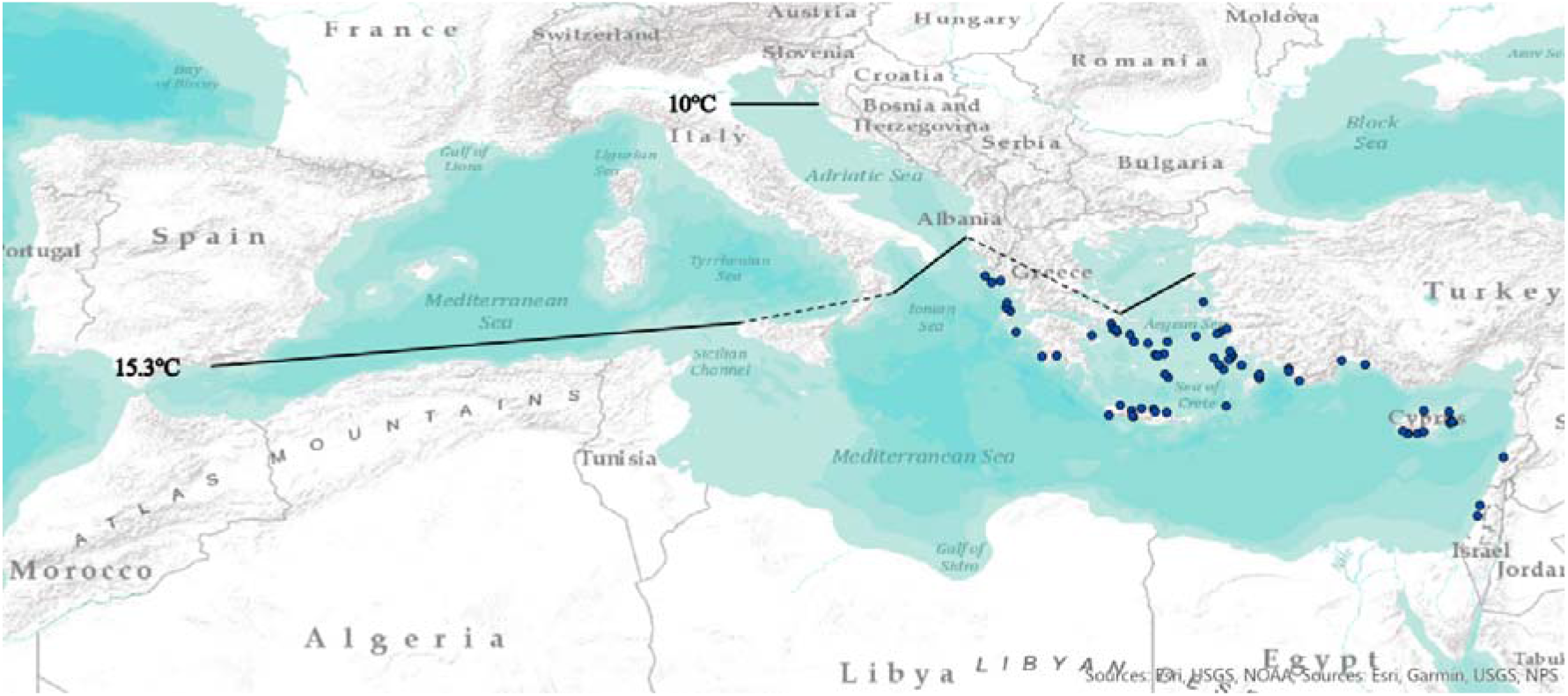
Thermal boundaries to invasion expansion. Black lines represent the thermal boundaries at 15.3°C and 10°C (as recorded by Dimitriadis et al., 2020; Johnston & Purkis, 2014) hypothesized to restrict lionfish expansion. Blue dots represent the lionfish sightings reported in our survey.

Behavioral flexibility and/or better cognitive abilities may be key to lionfish invasive success. Behavioral adaptations have been shown to increase success of invasive species across a variety of taxa (Weis & Sol, 2016). For instance, in round gobies (*Neogobius melanostomus*), higher dispersal potential and bolder personalities are shown in individuals on the invasive front, while in the common myna (*Acridotheres tristis*) invasive populations have increased innovation and decreased neophobia compared to native populations (Magory, Kumar, Nair, Hauber, & Dor, 2020; Myles-Gonzalez, Burness, Yavno, Rooke, & Fox, 2015). We predict that lionfish in invasive territories may similarly possess different behavioral and/or cognitive traits that are more advantageous to invasive ability compared to their native counterparts. For example, post-settlement invasive lionfish may find themselves in novel environments, so better operant learning or innovative ability might help these individuals adapt, survive, and establish in new areas (Weis & Sol, 2016). In this way, habitats or environments previously thought unsuitable for lionfish might become inhabited by invasive lionfish in the upcoming years. How invasive populations in the Atlantic and the Mediterranean differ in their behavior and cognition also has yet to be investigated, as this could cause differential expansion rate and invasive ability between the two populations.

Differences in predation level by filter feeders could also impact the precision of models in predicting invasive ability. The Enemy Release Hypothesis predicts that invaders escape population controls (predators, competitors, parasites, pathogens, etc.) present in their native range, causing them to thrive in novel invasive ecosystems (Mack et al., 2000; Torchin & Mitchell, 2004). Although adult and juvenile lionfish are found to have no natural predators in either their native or invasive ranges due to their venomous spines, larvae predation during the pelagic phase by filter feeders has yet to be explored (Côté et al., 2013; Galloway & Porter, 2019). Filter feeders in the native range may have evolved with lionfish to be able to feed on them as larvae, but those in invasive ranges have not, causing the higher population density of invasive lionfish compared to native populations, and further range expansion of these species then previously predicted (Darling, Green, O’Leary, & Côté, 2011). For example, a prevalent filter feeder in the Indo-Pacific, *Herdmania momus*, is also invasive in the Mediterranean via introduction through the Suez Canal (Rius & Shenkar, 2012). *H. momus’s* invasive range does not expand as far north or west as the lionfish’s however, suggesting that lionfish may have been able to spread far enough in the Mediterranean to escape the predation pressure of native filter feeders, allowing further expansion than predicted models (Çinar, Bilecenoglu, Öztürk, & Can, 2006; Gewing, Goldstein, Buba, & Shenkar, 2019). The absence of coral reefs in the Mediterranean Sea may also impact the predation pressure on lionfish larvae, as coral reef sponges are also known to decrease the abundance of pelagic larvae through filtration (Riisgård & Larsen, 2010).

Another question that remains is why the Mediterranean invasion is starting now, rather than at the opening of the Suez Canal in 1869. One reason may be that the Suez Canal was recently expanded in 2015, allowing more opportunity for passage from native habitats in the Red Sea to the Mediterranean, but this may not be the only reason. For example, global warming in previous years may have led to an increased opportunity for lionfish to establish and thrive in the Mediterranean (Gattuso et al., 2015; Hoegh-Guldberg & Bruno, 2010). Lionfish may have also developed a mutation causing better physiological cold resistance in invasive populations and the ability to reside closer to the temperature barrier. Additionally, adaptations that effect behavior could have made lionfish more able to adapt to Mediterranean conditions, similar to the blue tit innovation in the United Kingdom after urbanization in the 1940s. These birds learned to pierce the foil on the top of milk bottles as a novel way to obtain food in an increasingly urbanizing environment (Aplin, Sheldon, & Morand-Ferron, 2013). Similarly, lionfish may have learned and developed behavioral adaptations that have made them more likely to fit into an unoccupied ecological niche present in the Mediterranean, allowing them to thrive in the new niche (Dimitris Poursanidis et al., 2020). These types of adaptations in invasive populations should be tested to better understand the invasive ability of various species.

## Conclusions

Based on local dive center expertise we found that lionfish have spread and established in the eastern part of the Mediterranean as far as the Greek Ionian coast. We also found clear evidence of ongoing expansion with new sightings as recent as October 2020 (23.1537875°E, 37.6378821°N), showing the lionfish are still in the middle of their invasion. This expansion into new invasive territory demonstrates the need for continuous monitoring of this species using local expertise and development of management methods to protect native biodiversity (B. Galil, Marchini, Occhipinti-Ambrogi, & Ojaveer, 2017). For instance, careful documentation of lionfish distribution in the Caribbean has allowed for evaluation and implementation of management strategies (de León et al., 2013). We also argue that reliance on citizen may not tell the whole story, as it requires the general public to reliably identify invasive species, and for citizen science campaigns for certain species to be widely advertised in the area. For example, the sighting we found in Spain might have been unreported until this point because attention is not drawn to identifying and tracking lionfish there. Only local experts like those found at dive centers will notice the new species due to their familiarity with the area. Finally, to create effective management strategies for lionfish populations in the Mediterranean, we need to intensify our research effort to know more about why this species is able to expand its territory so quickly, even under unfavorable conditions.

